# Electrophysiological Dynamics of Default Network Suppression

**DOI:** 10.64898/2026.11.06.764967

**Authors:** Wendy Sun, Mark R. Libowitz, Rhiannon L. Cowan, Theo Tobel, Jingnan Du, Daniel A. Feldman, Tyler S. Davis, Jace B. King, Shervin Rahimpour, Elliot H. Smith, Randy L. Buckner, Ben Shofty

## Abstract

Distributed brain networks compete with one another in the service of information processing. One dramatic example is antagonism between networks that support internally directed cognition, often collectively referred to as the default network, and networks that orient to external events and drive action. Here we combined within-individual precision functional neuroimaging and intracranial electrophysiology to investigate network competition at high spatiotemporal resolution. In six patients, robust antagonism between networks was established during a target detection task. Network-assigned electrophysiological recordings displayed concordant responses, including in single-neuron firing rates, and further revealed the temporal dynamics of network competition. Networks involved in salience detection and action planning showed early, transient responses that peaked near the participant response. Default network suppression occurred later, reflected in decreases in high-frequency broadband activity and in single-neuron firing rates in response to targets. Examining subcortical regions revealed rapid activity increases in the thalamus and posterior hippocampus aligned to the participant response. Critically, a late evolving, sustained suppression was observed throughout the hippocampus. These findings suggest that, across cortical and subcortical regions, network competition is temporally layered. Furthermore, the hippocampus can be dissociated along its long axis during competition. Such push-pull antagonism may be fundamental to advanced human cognition, and its disruption may underlie mental illness.

## Introduction

Distributed brain networks that support cognitive function compete with one another during thought and action. These networks consist of association regions linked by long-range anatomical projections (Geschwind, 1965; Goldman-Rakic, 1988; Mesulam, 1990; 1998). A dramatic example of network competition emerges when participants shift between passive task states and states that demand active, attentive responses to external events (Gusnard and Raichle, 2001; Fransson, 2005; Fox et al., 2005; Anticevic et al., 2012; Buckner and DiNicola, 2019; Dosenbach, Raichle, and Gordon, 2025). Responding to external, salient stimuli robustly suppresses networks active during internal modes of cognition (often collectively referred to as the ‘default network’). This phenomenon was serendipitously discovered in the earliest brain imaging studies (Andreasen et al., 1995; Shulman et al., 1997; Raichle et al., 2001; Mazoyer et al., 2001) and has been the focus of extensive research in the decades since (for recent review see Buckner and DiNicola, 2019). Human (Miller, Weaver, and Ojemann, 2009; Ossandón et al., 2011; Dastjerdi et al., 2011; Foster, Dastjerdi, and Parvizi, 2012; Foster et al., 2015; see Fox et al., 2018 for review) and non-human primate (Hayden, Smith, and Platt, 2009) studies reliably reveal electrophysiological correlates of suppression when active, externally-oriented tasks are engaged. Moreover, the suppressive effect is temporally offset, with task-evoked activation preceding suppression (Raccah et al., 2018; Kucyi et al., 2020).

Candidate mechanisms of network competition linked to the default network have been suggested by two lines of evidence. The first line of evidence concerns network anatomy and function in the human brain. Networks that are suppressed preferentially associate with limbic structures including the hippocampus and amygdala (Shulman et al., 1997; Greicius et al., 2004; Vincent et al., 2006; Zheng et al., 2021; Reznik et al., 2023; Edmonds et al., 2024; Angeli et al., 2025). Furthermore, these networks support internal modes of cognition. For example, human functional connectivity from the hippocampus and adjacent parahippocampal cortex identifies a distributed cerebral network, referred to as Default Network-A (DN-A; Braga and Buckner, 2017; Braga et al., 2019), that is active when individuals mentalize scenes during remembering and imagining future events (DiNicola, Ariyo, and Buckner, 2023; Du et al., 2024; Angeli et al., 2025; Ladwig et al., 2026; see also Hassabis and Maguire, 2009). This network is suppressed during simple, stimulus-driven tasks (e.g., Du et al., 2024). A second, parallel network, referred to as Default Network-B (DN-B), is coupled to the entorhinal cortex and the amygdala (Reznik et al., 2023; Edmonds et al., 2024), and active during mentalizing tasks involving social inferences (DiNicola, Braga, and Buckner, 2020; Du et al., 2024; Deen and Freiwald, 2025; Ladwig et al., 2026). This second network is also robustly suppressed during the same stimulus-driven tasks (e.g., Du et al., 2024; see also Ladwig et al., 2026).

The second line of evidence comes from studies that have directly explored circuit-level mechanisms underlying network competition. Distributed cortical regions forming networks are integrated via thalamocortical circuits (Goldman-Rakic, 1988; Jones, 2009; Saalmann and Kastner, 2009; Zikopoulos and Barbas, 2012; Halassa and Kastner, 2017; Halassa and Sherman, 2019). Activity levels in the cortical networks are modulated by the thalamic reticular nucleus (TRN), which surrounds the thalamus proper (Pinault, 2004; Jones, 2007). Specifically, corticothalamic and thalamocortical axons send collaterals to the TRN, which in turn exerts control over information flow through GABAergic inhibition of thalamic projection neurons. Within the TRN are subpopulations of neurons linked to first-order (primary sensory) nuclei that can be distinguished from other subpopulations linked to second-order nuclei, including those associated with limbic structures (Li et al., 2020). Halassa and colleagues (2014) further demonstrated that TRN subnetworks can be mutually antagonistic: during sleep and rest, sensory-projecting TRN neurons show activity consistent with inhibition of sensory processing, whereas limbic-projecting neurons show the opposite effect.

These findings raise the possibility that the robust and ubiquitous human neuroimaging phenomenon of default network suppression arises from antagonism between networks that respond to immediate environmental events and hippocampal-associated networks that support internal modes of cognition. To resolve the dynamics of network competition, we combined precision functional MRI (fMRI) with intracranial electroencephalography (iEEG) recorded from depth electrodes capturing local field potentials and single-neuron firing. Critically, we measured neurophysiological suppression in the cortex as well as in the hippocampus. In addition, we recorded from the thalamus, given its role in integrating distributed cortical networks. Surgical patients were first scanned pre-operatively to estimate each individual’s idiosyncratic network organization with precision functional connectivity and confirm default network suppression using a target detection task. Following surgical implantation of macro-and micro-depth electrodes, we discovered a temporally layered structure of the antagonism. Rapid activity increases in the thalamus and posterior hippocampus, aligned to the participant response, were followed by a sustained, robust suppression of the hippocampus. This push-pull antagonism may reflect a fundamental dichotomy between external versus internal modes of cognition that, when in balance, supports the most advanced aspects of human cognition but, when disrupted, leads to mental illness, including psychosis.

## Results

### Target Detection Elicits Robust Antagonistic Suppression of the Default Network

Presurgical fMRI revealed robust antagonistic effects in each individual and across the group average (**Fig. 1**). Mapping the broad cortical effect revealed the canonical pattern that defines the default network: as networks involved in detecting and responding to the stimulus target increased their response (**Fig. 1**, white triangles), a broad relative reduction of the blood oxygenation level-dependent (BOLD) signal --defined here as suppression --was prominent across midline regions including medial prefrontal cortex (MPFC) and precuneus cortex, as well as lateral posterior parietal cortex (**Fig. 1**, white asterisks). This pattern of task-induced suppression, observed here within individuals, is remarkably similar to the pattern first identified in early human neuroimaging studies (e.g., Shulman et al., 1997; Mazoyer et al., 2001; Raichle et al., 2001). Additional activation was observed along the central sulcus in the left hemisphere near the hand representation, consistent with the right-handed button press, and along the calcarine sulcus near the central visual field representation, consistent with the target being a centrally-presented visual stimulus.

**Figure 1.**
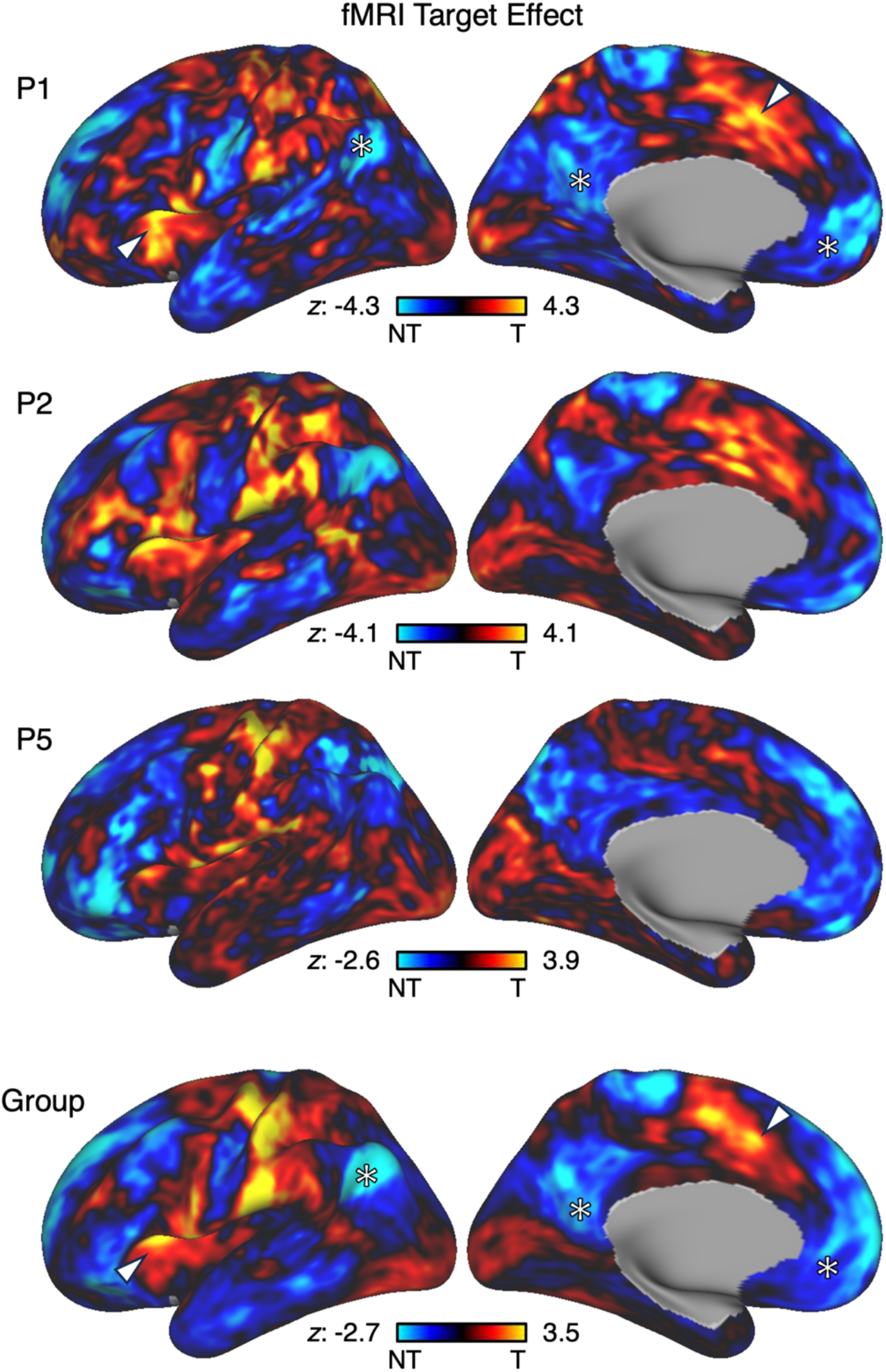
Target detection elicits opposing fMRI responses with suppression of regions canonically linked to the default network. The main task contrast (Target Effect; oddball targets > distractor nontargets) is shown on an inflated cortical surface, with z-statistical maps representing increases in red/yellow and decreases in blue. Data are unthresholded to allow visualization of the full effect in both directions. Maps are provided for three representative individuals (**Top Three Rows**) as well as for the average group of participants (**Bottom Row**). While idiosyncratic details differ by participant, the fMRI Target Effect yields activation across distributed regions including the anterior insula and dorsal anterior cingulate cortex (white triangles). By contrast, there is suppression across distributed regions typically associated with the default network, including medial prefrontal cortex (MPFC), precuneus cortex, and lateral posterior parietal cortex (white asterisks). T, targets; NT, nontargets.

Quantification of the push-pull response in the *a priori* within-individual defined networks revealed the antagonistic effect. The fMRI hemodynamic response during target detection was quantified for the two networks predicted to increase their activity (the salience network, SN, and the action-mode network, AMN; Seeley et al., 2007; Menon and Uddin, 2010; Seeley, 2019; Du et al., 2024; Dosenbach, Raichle, and Gordon, 2025), and separately the two networks hypothesized to decrease their activity (DN-A and DN-B). For the main analyses, AMN/SN were grouped and DN-A/DN-B were grouped for technical reasons, as the networks within each group are highly interdigitated and in the context of antagonism during target detection, functionally similar. A clear antagonistic response was observed in all six participants (**Fig. 2**, Left; see also Supplementary Fig. S2 for results in each network separately, ungrouped). Given the limited temporal resolution of the indirect neurovascular (BOLD) signal that fMRI detects, the relative timing of the antagonistic responses could not be resolved.

**Figure 2.**
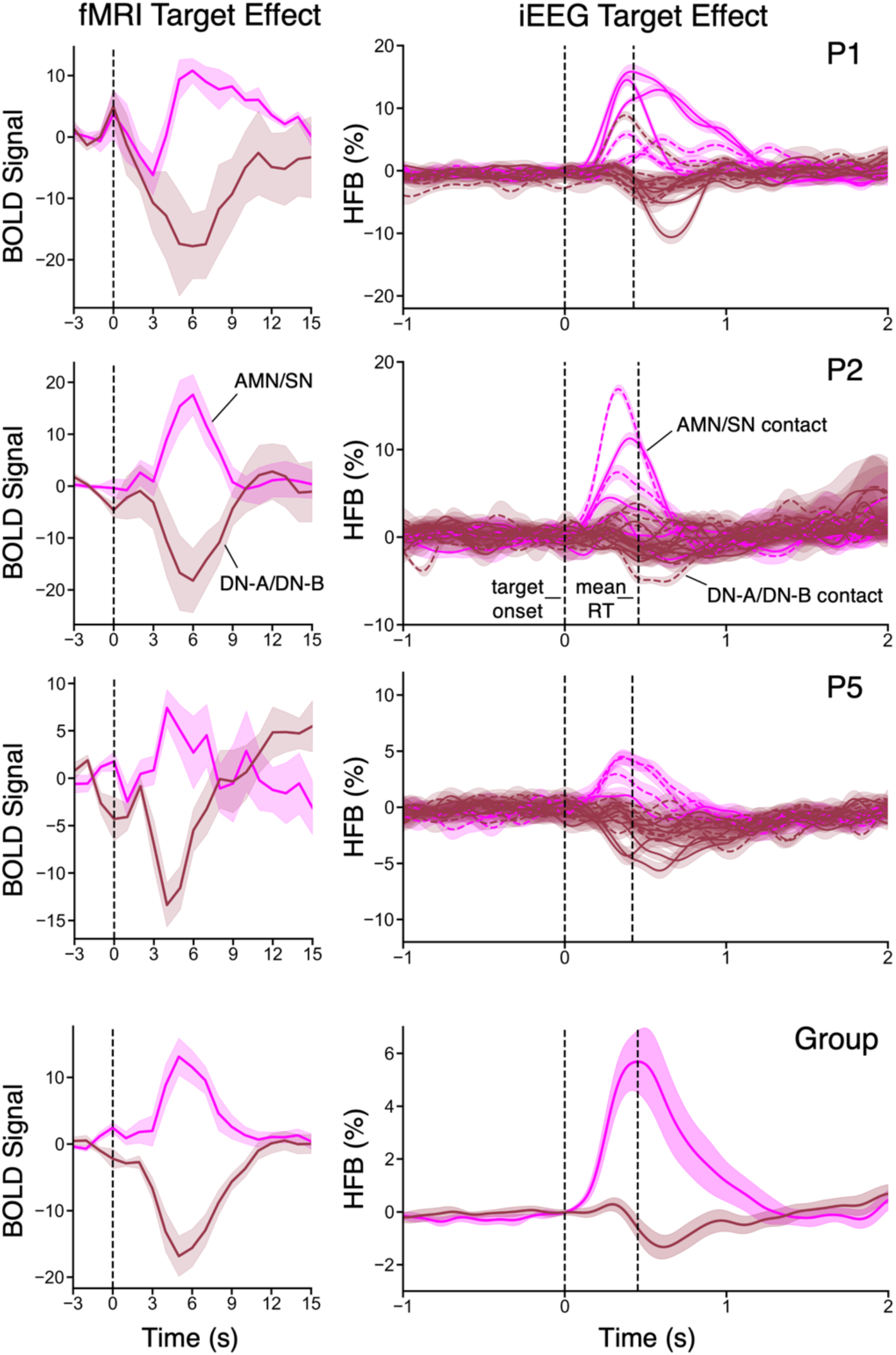
Functional network estimates predict iEEG activity in cortical contacts. The mean timeseries locked to the onset of oddball target trials, as contrast to distractor nontarget trials, is displayed for network-level BOLD activity within AMN/SN and DN-A/DN-B (**Left**), as well as for the corresponding HFB activity in all available cortical contacts localized to AMN/SN and DN-A/DN-B (**Right**). Timeseries plots are provided for three representative individuals (**Top Three Rows**) and for the average group of participants (**Bottom Row**). In the BOLD timeseries, solid curves show the mean across runs/participants, shaded ribbons mark the standard error of the mean at each timepoint, and vertical dashed lines at 0 represent target onset. The HFB timeseries follow the same convention, with dashed curves denoting contacts within white matter, solid curves denoting contacts within gray matter, and an additional vertical dashed line representing the mean reaction time. The fMRI Target Effect elicits a robust increase in BOLD activity within AMN/SN and a robust decrease within DN-A/DN-B. The iEEG Target Effect reveals HFB increases in AMN/SN contacts and decreases in DN-A/DN-B contacts. AMN/SN, Action-Mode Network / Salience Network; DN-A/DN-B, Default Network-A / Default Network-B.

### Cortical Network Estimates Predict iEEG High-Frequency Broadband Activity

fMRI-derived network estimates predicted the direction of high-frequency broadband (HFB) responses in cortical contacts (**Fig. 2**, Right). Contacts localized to AMN/SN showed HFB increases following target onset (relative to nontargets), whereas contacts localized to DN-A/DN-B showed HFB decreases (relative to nontargets). This correspondence was present in individual participants and in the group average. We define suppression here as a significant decrease in HFB activity in response to targets relative to nontargets (cluster-corrected *p* < 0.05). Across participants, 71% (17/24) of contacts localized to AMN/SN showed significant increases during the HFB trial epoch following target onset, as opposed to 8% (8/102) of contacts localized to DN-A/DN-B (Supplementary Fig. S3). By contrast, 0% of contacts localized to AMN/SN (0/24) showed significant decreases, compared to 24% of contacts localized to DN-A/DN-B (24/102).

In addition, HFB activity revealed a temporal feature that the BOLD signal could not resolve. At the temporal resolution of fMRI, the AMN/SN increase and DN-A/DN-B decrease appear simultaneous beginning about 2 sec after stimulus onset and peaking at about 6 sec (**Fig. 2**, Left). By contrast, the millisecond resolution of iEEG revealed that HFB decreases in DN-A/DN-B contacts tended to follow increases in AMN/SN contacts (**Fig. 2**, Right). This apparent temporal offset is quantified below and replicates prior work leveraging group-level atlas-based approximations of the salience and default networks (Kucyi et al., 2020).

### iEEG Target Effect Reveals Rapid HFB Increase in Thalamus and Sustained Hippocampal Suppression

We next examined task-evoked iEEG responses in the thalamus and hippocampus. Contact-level data from the first participant are shown in **Fig. 3**, with similar plots for all other participants provided in Supplementary Figs. S6-S14. Thalamic contacts displayed a rapid increase in HFB activity following target onset that appeared temporally coincident with the cortical AMN/SN response (**Fig. 3A**, Middle). Non-parametric cluster permutation tests confirmed statistically significant HFB increases in thalamic contacts, with a timing profile similar to that of the cortical AMN/SN response (**Fig. 3B**). Hippocampal contacts, by contrast, displayed a sustained late suppression that peaked after both the thalamic and the cortical AMN/SN increase (**Fig. 3A**, Bottom). This suppression is consistent with the hippocampus’ established connectivity to the default network (Greicius et al., 2004; Vincent et al., 2006; Zheng et al., 2021; Reznik et al., 2023; Edmonds et al., 2024; Angeli et al., 2025). Notably, hippocampal contacts were suppressed to a greater extent than cortical DN-A/DN-B contacts (compare **Fig. 3A** Bottom to Top), distinguishing the hippocampal response in both magnitude and timing.

**Figure 3.**
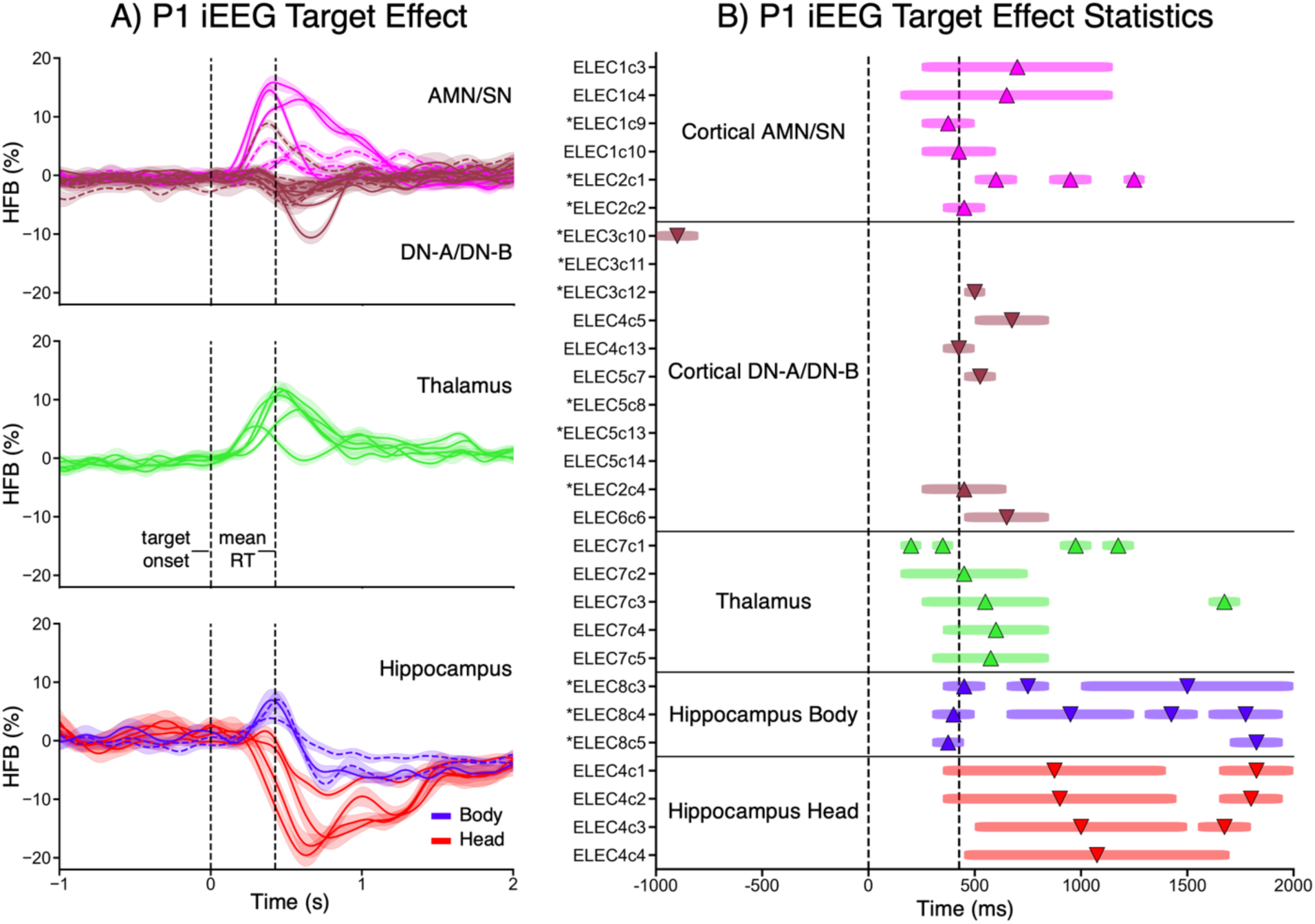
Task-evoked network competition extends to the thalamus and hippocampus. (**A**) The mean HFB timeseries anchored to the onset of oddball target trials, relative to distractor nontarget trials, is displayed for all available contacts localized to cortical AMN/SN and DN-A/DN-B, thalamus, and hippocampus in the first participant. Dashed vs solid curves denote contacts within white vs gray matter, and shaded ribbons mark the standard error of the mean at each timepoint. Vertical dashed lines mark target onset (0 sec) and mean reaction time. In addition to cortical network effects, there is a rapid activity increase in thalamic contacts, followed by a sustained late suppression in hippocampal contacts. Contacts localized to the anterior hippocampus (head) showed only late suppression, while those in the posterior hippocampus (body) showed an early transient increase, followed by late suppression. These network competition effects are supported by formal statistical testing. (**B**) Results of non-parametric cluster permutation tests assessing whether each contact’s HFB contrast differed significantly from zero (cluster-corrected *p* < 0.05). Contacts are grouped and colored by network/region. Horizontal colored bars indicate the time span of statistically significant clusters; each bar spans one or more contiguous 50 ms bins. Upward triangles denote significant HFB increases; downward triangles denote significant HFB decreases. An asterisk (*) preceding a contact label denotes a white matter contact. AMN/SN, Action-Mode Network / Salience Network; DN-A/DN-B, Default Network-A / Default Network-B.

Within the hippocampus, anterior (head) and posterior (body) contacts exhibited distinct response profiles (**Fig. 3A**, Bottom). In the first participant, contacts in the hippocampal body displayed a biphasic response consisting of an early transient activation and late sustained suppression. By contrast, contacts in the hippocampal head showed only late suppression. Cluster-based permutation tests in this participant showed statistically significant early increases followed by late decreases in hippocampal body contacts and only late decreases in hippocampal head contacts (**Fig. 3B**). The pattern replicated in additional participants (Supplementary Fig. S4): 57% (16/28) of contacts localized to the hippocampal body showed significant increases during the HFB trial epoch following target onset, as opposed to 9% (4/44) of contacts localized to the hippocampal head. Contacts localized to both the hippocampal body (89%; 25/28) and hippocampal head (95%; 42/44) consistently showed significant decreases (93% overall; 67/72). Hippocampal effects persisted when seizure onset zone contacts were excluded (compare Supplementary Fig. S4B and S4C).

To provide a summary view, HFB timeseries were averaged across all available contacts localized to each network and subcortical region within individuals and for the group (Supplementary Fig. S5). Effect magnitudes varied across individuals, and some were noisier than others, but an overall temporal structure became evident with multiple replications across individuals and convergence at the group level.

### Network Competition Follows a Temporal Sequence with Cortical and Thalamic Activation Preceding Sustained Hippocampal Suppression

Three metrics were used to quantify the temporal sequence observed in the HFB timeseries: onset latency, time-to-extremum (peak/trough), and cross-correlation lag. Of 133 contacts showing statistically significant HFB increases or decreases beginning within 1000 ms of target onset, 126 (95%) responded in the direction predicted by their network or regional assignment. These conforming contacts were included in the primary analysis (see Methods for exact criteria). Five of the seven non-conforming contacts were from a single participant (P6; Supplementary Table S3). However, the temporal ordering reported below was preserved when non-conforming contacts were included and when a direction-agnostic onset definition was applied (see Supplementary Table S4).

**Fig. 4** displays onset latency and time-to-extremum values in individual participants as well as the group (plots for all participants in Supplementary Fig. S15). On average, descriptively, cortical AMN/SN activated to targets first (onset latency 225 ± 43 ms), followed by the hippocampal body (280 ± 27 ms) and the thalamus (313 ± 75 ms). Then, suppression began in cortical DN-A/DN-B (481 ± 47 ms), followed by hippocampal head (517 ± 68 ms) and body (625 ± 108 ms). For time-to-extremum, the same descriptive ordering was observed, except that the hippocampal body increase peaked before cortical AMN/SN. Hippocampal suppression reached its trough later than any other region’s time-to-extremum (head, 779 ± 202 ms; body, 921 ± 220 ms; Supplementary Table S5; see also Supplementary Fig. S15). Within hippocampal body contacts, the interval from transient increase (peak) to late suppression (trough) had a median of 532 ms (IQR = 372–728 ms; range = 222–1364 ms), supporting a biphasic response profile and providing a candidate temporal account of the posterior hippocampal transients reported by Angeli et al. (2025).

**Figure 4.**
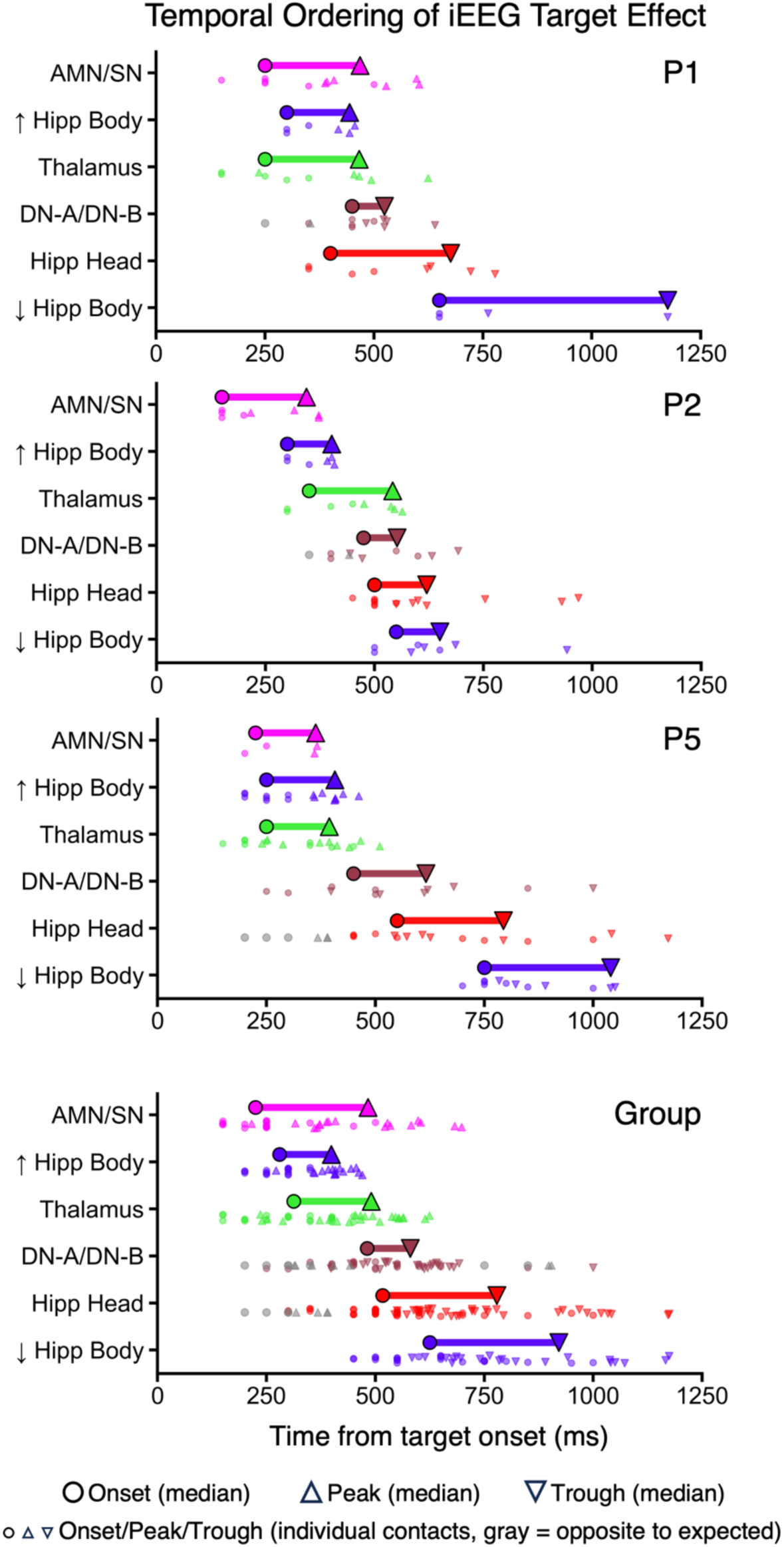
Thalamic and cortical activation precedes hippocampal suppression in a consistent temporal sequence. The onset latency and time-to-extremum of the task-evoked HFB response, locked to oddball target onset, are summarized in three representative individuals (**Top Three Rows**) and for the group average (**Bottom Row**). For each network/region, a shaded bar spans the median onset (shaded circle) to the median time-to-extremum (shaded triangle, pointing upward for peaks and downward for troughs). The group panel reflects average values across all available participants. Smaller markers below each bar show values for individual contacts. Gray markers indicate contacts with significant deflections opposite to the expected direction; these are plotted at their observed onset and extremum but did not contribute to primary timing estimates. In the group average, AMN/SN and thalamic increases preceded DN-A/DN-B and hippocampal decreases, with the hippocampal body trough occurring last. AMN/SN, Action-Mode Network / Salience Network; DN-A/DN-B, Default Network-A / Default Network-B; Hipp, Hippocampus.

Cross-correlation analysis was used to quantify the temporal offset between pairs of network-and region-averaged HFB waveforms. At peak negative correlation (r = −0.75), AMN/SN led cortical DN-A/DN-B by 181 ± 66 ms. AMN/SN and thalamic responses were positively correlated (r = 0.87) and approximately synchronous; AMN/SN and hippocampal body increases were also positively correlated (r = 0.65) and approximately synchronous. Cortical DN-A/DN-B and hippocampal head suppression were positively correlated (r = 0.68, with cortical suppression leading hippocampal head suppression by 133 ± 114 ms). All computed pairwise temporal relationships are provided in Supplementary Table S6.

### The Temporal Sequence of Network Competition Spans the High-Frequency Spectrum

To characterize network competition across the full frequency range, time-frequency spectrograms (3–250 Hz) were computed for cortical network contacts (**Fig. 5**) and for thalamic and hippocampal contacts (**Fig. 6**). In cortical AMN/SN contacts, target onset elicited broadband amplitude increases in the gamma and high-gamma ranges (∼30 Hz and above), including the 75–175 Hz high-frequency band visualized in **Fig. 2**. In cortical DN-A/DN-B contacts, the same frequency ranges showed amplitude decreases. Effects below the gamma range (< ∼30 Hz) are displayed for completeness but not interpreted. As in the HFB timeseries, the temporal offset between AMN/SN activation and DN-A/DN-B suppression is apparent in the spectrograms (**Fig. 5**).

**Figure 5.**
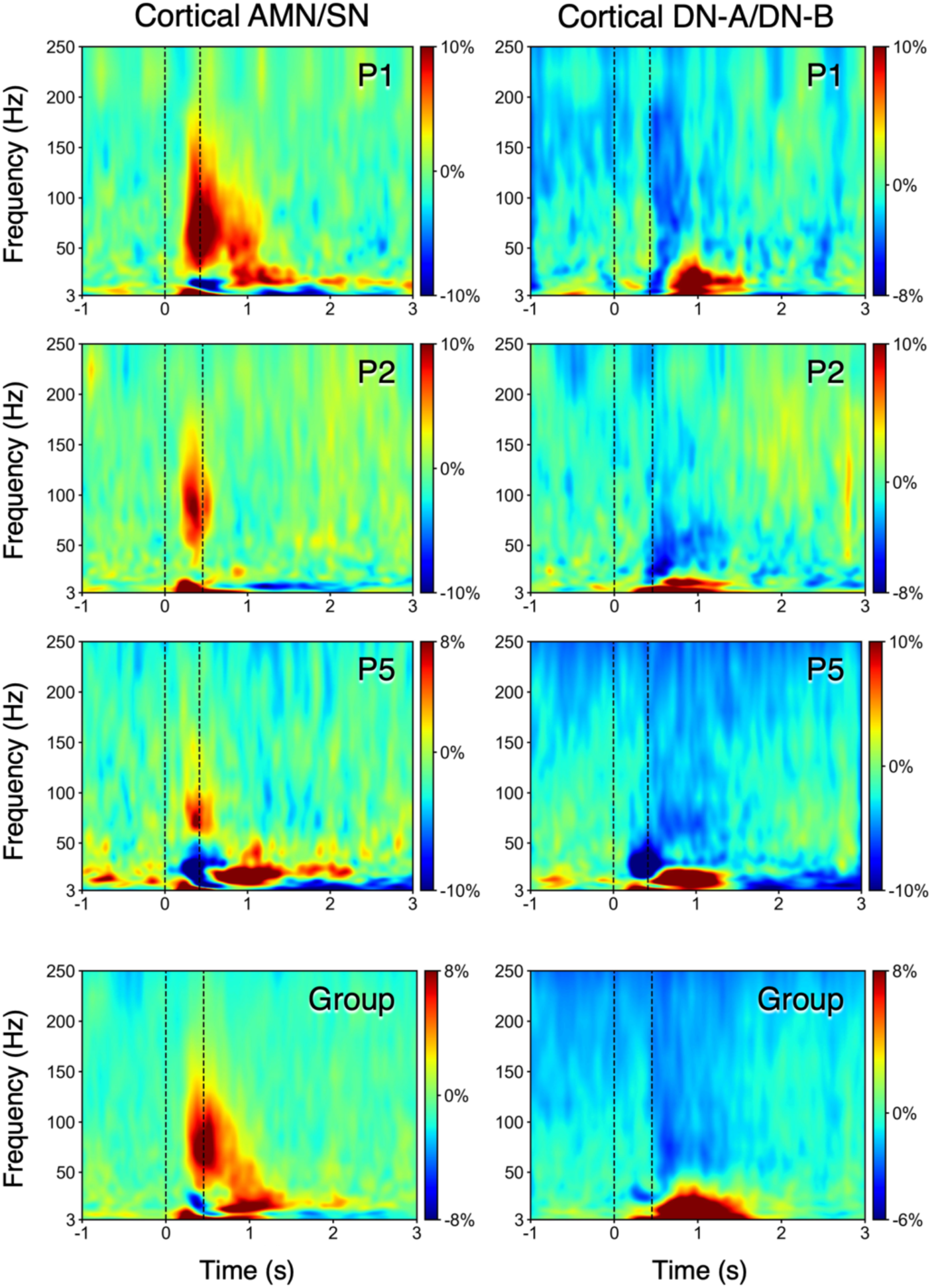
Mean time-frequency spectrograms illustrate opposing effects in cortical network contacts. Time-frequency spectrograms (3-250 Hz) are shown in three representative individuals (**Top Three Rows**) as well as in the average group of participants (**Bottom Row**). Spectrograms reflect % change in amplitude envelope of oddball target trials relative to distractor nontarget trials, with increases in red/yellow and decreases in blue. Vertical dashed lines mark target onset (0 sec) and mean reaction time. Each plot takes the mean across all available contacts/participants for cortical AMN/SN and DN-A/DN-B, respectively. Across gamma and high gamma frequency ranges (∼30 Hz and above), AMN/SN contacts exhibit early increases, while DN-A/DN-B contacts exhibit later decreases. AMN/SN, Action-Mode Network / Salience Network; DN-A/DN-B, Default Network-A / Default Network-B.

**Figure 6.**
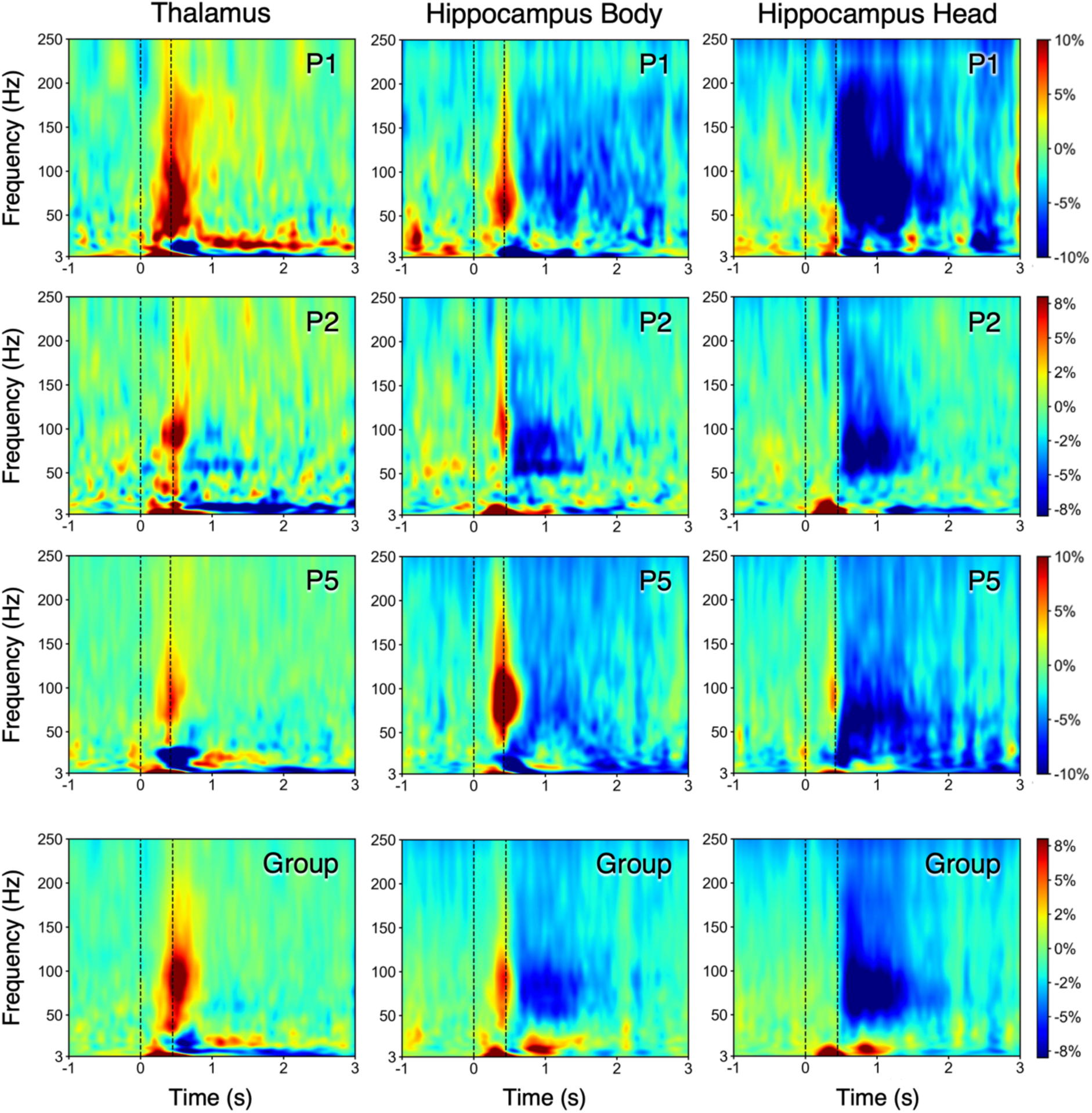
Mean time-frequency spectrograms illustrate thalamic activity increase and biphasic hippocampal response. Time-frequency spectrograms (3-250 Hz) are shown in three representative individuals (**Top Three Rows**) as well as in the average group of participants (**Bottom Row**). Spectrograms reflect % change in amplitude envelope of oddball target trials relative to distractor nontarget trials, with increases in red/yellow and decreases in blue. Vertical dashed lines mark target onset (0 sec) and mean reaction time. Each plot takes the mean within all available contacts/participants for the thalamus, hippocampal body, and hippocampal head, respectively. In the gamma and high-gamma ranges (∼30 Hz and above), thalamic contacts show an early activity increase. Hippocampal body contacts exhibit an early activity increase followed by a late sustained decrease, a biphasic profile that is largely absent in hippocampal head contacts, which show a late sustained decrease.

Thalamic contacts showed broadband amplitude increases in the gamma and high-gamma ranges following target onset (**Fig. 6**). Hippocampal body contacts displayed an early transient increase followed by a late decrease, mirroring the biphasic HFB profile observed in **Fig. 3**. Hippocampal head contacts showed sustained amplitude decreases. These spectral patterns were present in individuals and in the group (**Figs. 5** and **6**, Bottom Rows).

### Putative AMN/SN and DN-A/DN-B Principal Cells Show Opposing Responses to Targets

Finally, we examined firing rates (FR) of single neurons from microwires that localized to distinct personalized networks, sorting these cells into putative cell types based on their action potential waveform features. Putative principal cells localized to AMN/SN (n = 57) showed greater firing in response to target versus nontarget trials (two-sided sign-flip permutation test, ΔzFR = 0.09, *p* = 0.006 -0.03) from approximately 0.30 -0.40 s (**Fig. 7E**). Nontarget trial firing remained near baseline (mean zFR = −0.008), whereas target firing averaged 0.09 zFR near 0.35 s (peak).

**Figure 7.**
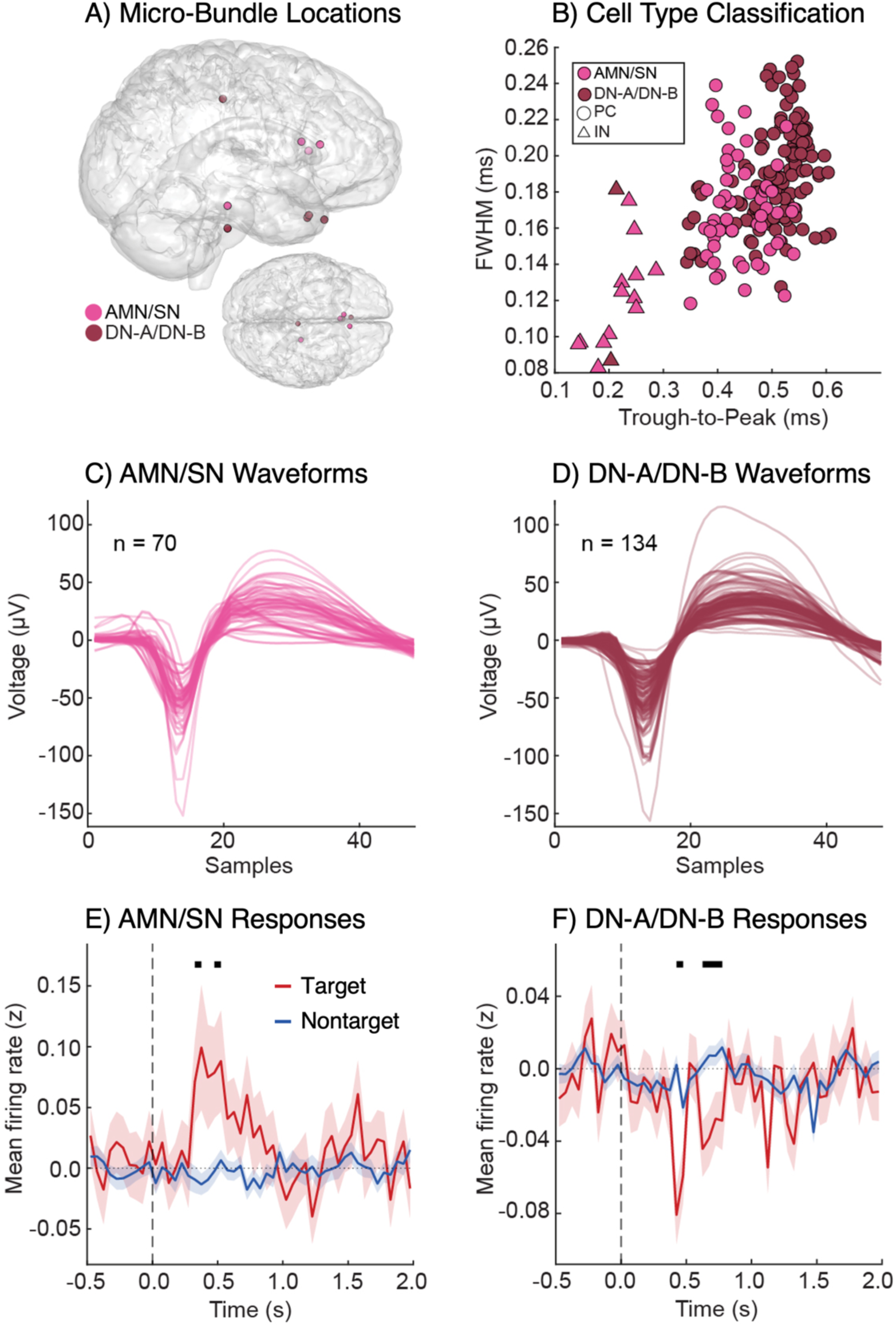
Opposing single-neuron responses to targets. (**A**) Anatomical locations of microelectrode bundles (n = 9 total; 4 AMN/SN and 5 DN-A/DN-B) across three patients. (**B**) Scatter plot of putative cell type classification using waveform shape metrics: full-width-at-half-maximum (FWHM; ms) and trough-to-peak (ms) durations. Triangles represent putative fast-spiking interneurons; circles represent putative principal cells (total isolated neurons = 204). (**C**) Isolated waveforms from AMN/SN (n = 70). Samples represent sampling rate (30 kHz) by time (1 sample = 33.33 μs). (**D**) Isolated waveforms from DN-A/DN-B (n = 134). (**E**) Population responses of putative AMN/SN neurons to targets. Mean z-scored firing rates significantly increase in response to targets (red trace) compared to nontargets (blue trace). Horizontal black bars indicate time bins with significant target-nontarget differences. (**F**) Population responses of putative DN-A/DN-B neurons to targets. Mean z-scored firing rates significantly decrease in response to targets (red trace) compared to nontargets (blue trace). Horizontal black bars indicate time bins with significant target-nontarget differences. AMN/SN, Action-Mode Network / Salience Network; DN-A/DN-B, Default Network-A / Default Network-B; PC, Principal Cell; IN, Interneuron.

Putative principal cells localized to DN-A/DN-B (n = 132) showed reduced firing in response to target relative to nontarget trials (two-sided sign-flip permutation test, ΔzFR = −0.06, *p* = 0.001 -0.04) from approximately 0.40 -0.50 s, with a second significant window from approximately 0.60 -0.80 s (ΔzFR = −0.04, *p* = 0.005 -0.03; **Fig. 7F**). Nontarget firing remained near baseline during these periods (mean zFR = −0.009 and 0.007), while target firing reached its strongest window-averaged decrease of −0.07 zFR near 0.45 s (trough), later than the AMN/SN peak. Single-neuron firing thus mirrored the direction of the HFB effects in each network.

## Discussion

Dynamic competition between networks is a key feature of brain function. By combining individualized precision neuroimaging with intracranial recordings, we discovered that network competition has a consistent temporal signature. Early activity evolving in regions driving task performance occurs along with a rapid increase in thalamic activity. Suppression of the default network follows, including a robust, sustained suppression of activity in the hippocampus, which was observed in nearly all hippocampal contacts. The hippocampal response further dissociates along the long axis, with an early positive transient response in the posterior hippocampus that is directionally and temporally distinct from a robust, broad suppression observed throughout the hippocampus. Below, we consider circuit mechanisms that may give rise to these dynamics.

### Action-Mode and Salience Activation Precede Default Network Suppression in Cortical Contacts

During target detection, opposing network responses were observed in the BOLD signal, consistent with expectations from prior work (e.g., compare **Fig. 1** to Du et al., 2024 Fig. 26). The same opposition was recapitulated in the electrophysiological response within individuals: following target onset, cortical contacts in the action-mode and salience networks showed increases in HFB activity, while contacts in the default network showed decreases (**Fig. 2**), supported by formal statistical testing (**Fig. 3**). Time-frequency spectrograms revealed gamma and high-gamma (∼30 Hz and above) amplitude increases in AMN/SN contacts and decreases in DN-A/DN-B contacts (**Fig. 5**), suggesting that the network competition effect is not limited to broadband averaging. Concordant evidence from our most direct measure, single-neuron recordings of isolated principal cells, showed increased firing rates in putative AMN/SN neurons and reduced firing rates in putative DN-A/DN-B neurons during target detection, supported by formal statistical testing (**Fig. 7**).

Importantly, the millisecond resolution of iEEG revealed that the AMN/SN activation and DN-A/DN-B suppression were not simultaneous: increases in AMN/SN contacts preceded decreases in DN-A/DN-B contacts (**Figs. 3** and **4**). Single-neuron firing rate effects also appear temporally offset, with the trough in DN-A/DN-B occurring after the peak in AMN/SN (**Fig. 7**). This ordering converges with prior intracranial findings that anatomical/atlas-based approximations of externally oriented networks lead default network suppression, reported for the dorsal attention network during arithmetic processing (Raccah et al., 2018) and for both dorsal attention and salience networks during withholding to rare targets (Kucyi et al., 2020). Our findings reinforce and extend those observations. The effects reported here emerge from within-individual networks defined *a priori*, demonstrating that precision estimates from a single, low-burden fMRI session (∼1 hour) can anticipate the direction of the HFB response at the level of individual contacts, consistent with and extending atlas-based localization approaches (compare Supplementary Fig. S3C to Kucyi et al., 2020 Fig. 1e).

### The Hippocampus Dissociates Along Its Long Axis During Network Competition

Hippocampal activity dynamics were not uniform along its long axis. Contacts in the anterior hippocampus (head) tended to show only late suppression, whereas contacts in the posterior hippocampus (body) tended to show a biphasic response: an early transient HFB increase, time-locked to the target and overlapping with cortical AMN/SN and thalamic activations, followed by the same late suppression seen in anterior contacts (**Figs. 3, 4, 6**). Task-evoked hippocampal responses to rare target events have been documented since early intracranial and magnetoencephalographic recordings (McCarthy et al., 1989; Tesche, Karhu, and Tissari, 1996; Kukleta et al., 2003; reviewed in Halgren, Marinkovic, and Chauvel, 1998), and the hippocampus has been implicated in novelty detection (Knight, 1996; Vinogradova, 2001; Kumaran and Maguire, 2009).

Recent precision functional neuroimaging work has situated this target/novelty response along the long axis and related it to hippocampal participation in distinct cortical networks (Angeli et al., 2025). Our results suggest an electrophysiological account for these observations. Although the hippocampus is correlated with the default network along its full extent, its anterior and posterior components preferentially couple to internally and externally oriented networks, respectively, with the posterior hippocampus tracking oddball targets and task transitions. The present results reveal an early transient response in the posterior hippocampus consistent with this target-related response (**Fig. 6**).

What is most striking is that, following the initial rapid activity increase, a robust and sustained suppression of activity in the hippocampus emerged. The effect was evident in the HFB timecourses of individual participants (**Fig. 3**) as well as in the timing estimates for individuals and the group (**Fig. 4**). Additionally, the hippocampal suppression was robust in the time-frequency spectrograms and distinct from the early transient increase in HFB activity (**Fig. 6**).

### Considerations for Thalamic Control of Network Competition

Cortical network activation was accompanied by a robust response in thalamic contacts. Following target onset, thalamic contacts showed a rapid increase in HFB activity that was temporally aligned with the cortical AMN/SN increase and preceded DN-A/DN-B and hippocampal suppression (**Figs. 3** and **4**). This response was consistent: 92% (22/24) of thalamic contacts showed a significant HFB increase, and the effect was also evident in the time-frequency domain as a broadband increase in the gamma and high-gamma ranges (**Fig. 6**). Although we cannot resolve which network(s) these thalamic contacts are related to, it is useful to hypothesize potential circuit implications.

The thalamic activity increase preceded the hippocampal suppression, a temporal pattern that constrains potential mechanisms of thalamocortical control. One candidate circuit involves the TRN, which surrounds the thalamus proper and exerts its influence through GABAergic inhibition of thalamic projection neurons (Pinault, 2004). Optogenetic work in mice has shown that the TRN comprises sensory-and limbic-projecting subnetworks with opposing, state-dependent effects: the sensory subnetwork disengages during external attention, while the limbic subnetwork is most quiescent during low-arousal states (Halassa et al., 2014). These circuit observations offer a potential mechanism by which thalamic activity could relate to the robust hippocampal suppression observed here. The hippocampal formation is connected with posterior cingulate and retrosplenial cortex (Kobayashi and Amaral, 2003; 2007), which in turn connect with the anterior thalamic nuclei (Vogt, Rosene, and Pandya, 1979) gated by the limbic-projecting TRN subnetwork (Halassa et al., 2014).

The observed temporal order of changes in the brain, with cortical DN-A/DN-B suppression preceding the robust, sustained hippocampal suppression, is consistent with the hippocampus being modulated through this thalamocortical circuit. We emphasize, however, that this account is a reasoned hypothesis. Our HFB recordings from clinical macrocontacts cannot distinguish reticular from relay neurons or resolve which exact thalamic territory the activity increase arises from, leaving the source of the thalamic contribution unresolved. Establishing whether the thalamus shapes the hippocampal response will require causal perturbation during simultaneous recording.

### Limitations and Future Directions

Electrode placement was determined entirely by clinical considerations, resulting in unevenly distributed anatomical coverage, with contacts concentrated in lateral temporal and frontal cortices (Supplementary Fig. S3A, B). As such, multiple cortical regions associated with the default network (e.g., posterior cingulate / retrosplenial cortex, inferior parietal lobule) were not well sampled in the present study and this may be reflected in the observation that cortical default network suppression was qualitatively the weakest magnitude effect (in contrast to prior studies, e.g., Fox et al., 2018). The hippocampus, by contrast, was densely sampled in the present study and revealed robust suppression. A larger cohort with more distributed coverage would help resolve whether the modest cortical default network effect reflects sparse and variable sampling or a genuine difference in response strength.

Another limitation is that we recorded from a cohort of patients with epilepsy, and many hippocampal contacts fell within clinically defined seizure onset zones (Supplementary Table S1). In contact-level visualizations and statistics, we explicitly label contacts residing within epileptologist-defined seizure onset zones. However, several factors mitigate the influence of epileptiform activity on our results. No patient experienced a seizure during testing, and we applied an empirical outlier-based rejection procedure to attenuate the contribution of interictal epileptiform discharges (IEDs). The hippocampal suppression and posterior biphasic response also persisted when all seizure onset zone contacts were excluded from the analysis (compare Supplementary Fig. S4B and S4C).

We have also treated HFB increases and decreases as markers of changes in local neuronal activity, a relationship established in cortex but less certain in the hippocampus, where coupling between HFB and the hemodynamic signal has been suggested to be weak or negative (Ekstrom et al., 2009; Hill et al., 2021). Our interpretation does not necessarily depend on this coupling: the link between HFB power and neuronal firing holds across regions (Manning et al., 2009), and the hippocampus is a robust and reproducible node of the default network (Greicius et al., 2004; Vincent et al., 2006; Kahn et al., 2008; Zheng et al., 2021; Reznik et al., 2023; Angeli et al., 2025), consistent with the suppression we observe. In future explorations, single-unit and laminar recordings could clarify how the presently observed broadband response relates to the firing of specific hippocampal neuronal populations.

Additionally, for single-neuron analyses, our spatial localization of the microwire recordings was done by visual inspection at consensus meetings. This is because fMRI-defined networks with resolution at the voxel level could not be unambiguously distinguished at the level of individual recording sites with resolution at the neuron level. We thus refer to the single-neuron localizations as putative. Importantly, localization was finalized prior to analysis.

Finally, our findings carry translational implications. The default network and its interactions with other large-scale networks are dysregulated in schizophrenia and in bipolar disorder with psychotic features (Whitfield-Gabrieli et al., 2009; Baker et al., 2014; Woodward and Heckers, 2016), conditions in which the boundary between internally generated and externally driven processing is thought to break down. Disrupted thalamocortical control may contribute to the network dysregulation seen in mental illness, in which case a paradigm that elicits and quantifies network antagonism directly could offer a circuit-level readout of that dysfunction within individual patients. By grounding network competition in its underlying electrophysiology, our findings are a step towards translating this measure into clinically relevant biomarkers, as well as providing direct physiological observations in humans that can be used to ground mechanistic studies in mouse and primate models.

## Conclusion

We employed direct electrical recordings from the human brain to discover a temporally layered sequence of brain network competition. Early corticothalamic activity temporally aligned to the active components of the task precedes suppression of the default network, prominently including the hippocampus. Within the hippocampus, posterior portions briefly increased activity before a broad suppression emerged along the entire sampled extent of the hippocampal long axis. Such dynamics suggest human network competition arises from broad opposition between cortical networks involved in goal-directed behavior and those linked to the hippocampus that support internal mentation.

## Materials and Methods

### Overview

This study combined precision functional neuroimaging with intracranial electrophysiology in neurosurgical patients to interrogate network competition. Supplementary Fig. S1 displays the experimental workflow. Pre-operatively, patients underwent an fMRI session that included a target detection task. From these data, individualized functional network estimates and task effects were extracted using an automated pipeline. Patients underwent clinical macroelectrode implantation and, post-operatively, performed the same target detection task during iEEG recording. Electrode contacts were localized to each patient’s functional networks and select subcortical regions for hypothesis-driven analysis.

### Participants

Adult patients (N = 6; mean age = 33.2 yr, SD = 5.7 yr; 2 female) undergoing intracranial monitoring for neurosurgical treatment of their drug-resistant epilepsy participated in a pre-operative fMRI scan and post-operative iEEG testing. Each participant provided informed consent through a protocol approved by the University of Utah Institutional Review Board. Participant eligibility criteria included age ≥ 18 yr, ability to provide informed consent, English as their first language, IQ ≥ 70, no history of a resective surgery, and no significant medical comorbidities (following Libowitz et al., 2026). Participants were labeled *P1*-*P6* and information about their demographics, implants, and seizure onset zone localization is available in Supplementary Table S1.

### Target Detection Task Paradigm

Participants were asked to complete a target detection task pre-operatively in the fMRI sessions and post-operatively during iEEG recordings. In this task, participants were instructed to respond to salient, visual targets that were uncommon relative to distractor nontargets (Du et al., 2024; Wynn et al., 2015). This task has been shown to elicit robust antagonism between networks involved in external orienting and those involved in internal mentation (Du et al., 2024). Participants each completed 5 runs of the target detection task in the fMRI session, and 10-14 runs in iEEG sessions. Overall behavioral performance was high, indicating task compliance (Supplementary Table S2).

Stimuli consisted of uppercase letters (O and K) presented in either black or red. Participants were instructed to press a button with their right index finger upon seeing a red K and to withhold responses to all other letter–color combinations. Trial composition within each run was 10% target red Ks, 10% lure red Os, 40% distractor black Ks, and 40% distractor black Os, with order randomized using Optseq (Dale, 1999). Each run lasted 350 sec (5 min 50 sec). Following 6 sec of fixation, the run began with a 20 sec fixation block, followed by a continuous sequence of 300 rapid event-related trials (each 1 sec: 0.15 sec letter presentation, 0.85 sec fixation), and ended with a final 20 sec fixation block. Start and end cues (1 sec “Begin” or “End,” followed by 1 sec fixation) bracketed the trial sequence.

### Pre-operative MRI Data Acquisition, Quality Control and Preprocessing

Pre-operative neuroimaging sessions were conducted at the University of Utah Imaging & Neurosciences Center using a 3T Siemens MAGNETOM Prisma^fit^ MRI scanner equipped with a 64-channel head coil (Siemens Healthineers AG, Erlangen, Germany). Structural images were acquired with the FreeSurfer recommended T1-weighted (T1w) multi-echo magnetization-prepared rapid acquisition gradient-echo (ME-MPRAGE) sequence: voxel size = 1.0 mm, TR = 2530 ms, TE = 1.69, 3.55, 5.41, and 7.27 ms, TI = 1100 ms, flip angle = 7°, matrix 256 × 256 × 192, 192 slices, in-plane GRAPPA acceleration = 2. As backup, a rapid T1w ME-MPRAGE (van der Kouwe et al., 2008) providing 1.2-mm isotropic voxels was acquired (TR = 2200 ms; TE = 1.57, 3.35, 5.13, 6.91 ms; TI = 1100 ms; flip angle = 7°; matrix = 210 × 210 × 192; in-plane GRAPPA acceleration factor = 4). Ten functional runs were acquired using a multiband gradient-echo echo-planar imaging (EPI) sequence sensitive to the blood oxygenation level-dependent (BOLD) contrast providing 2.4-mm isotropic voxels (TR = 1000 ms; TE = 33 ms; flip angle = 64°; matrix = 92 × 92 × 65). All BOLD runs began with 6 sec of fixation for T1 stabilization and lasted 350 sec. To mitigate spatial distortions, dual-gradient-echo B0 field maps were acquired with slice prescription and resolution matched to the BOLD sequence (TE = 4.45 and 6.91 ms). Participants viewed stimuli presented via PsychoPy software (Peirce et al., 2019) and projected to a display that was calibrated to ensure a central and comfortable view. Participants were instructed to stay still and alert prior to each scan.

Each structural scan and BOLD run was examined for quality. Images were manually inspected for incomplete head coverage, ringing, and ghosting. Automated quality control (QC) metrics were computed for every BOLD run, including slice-based signal-to-noise ratio (SNR) and maximum absolute motion. Qualitative raw data inspection and quantitative QC metrics were used to inform BOLD run exclusions, and all exclusions were finalized prior to analysis. In general, runs were excluded if SNR < 100 (borderline runs were examined case-by-case and kept if visual inspection deemed adequate quality). There were 9-10 usable runs (52-57 min of data) in each participant. Supplementary Table S2 provides data availability/inclusion details in each participant.

Data were preprocessed using the openly available “iProc” pipeline (see Braga et al., 2019 for in-depth description; Sun et al., 2025 for exact implementation details). The pipeline runs FreeSurfer *recon-all* (v6.0.0; Fischl, 2012) on the T1w image to generate a cortical ribbon estimate and automated anatomical segmentation including subcortical structures (“aparc+aseg”; Fischl et al., 2002; Desikan et al., 2006). To minimize spatial blur and preserve detailed functional anatomy within each individual, iProc combines all spatial transformations into a single interpolation applied to each raw BOLD volume: 1) alignment of all volumes in a BOLD run to the middle volume of that run, 2) B0 field map unwarping, 3) registration of all BOLD data to a mean BOLD template, and 4) registration of the mean BOLD template to the native-space T1w. All interpolated outputs were checked for registration errors. Following interpolation, nuisance variables (head motion parameters, whole-brain signal, ventricular signal, deep white matter signal, plus their temporal derivatives and quadratic terms) were regressed out, and high-motion volumes (framewise displacement > 0.4 mm) were censored. Data were then bandpass filtered at 0.01–0.1 Hz for network estimation only, and both filtered and unfiltered data were resampled to the fsaverage6 cortical surface mesh and smoothed with a 4-mm full-width-at-half-maximum (FWHM) Gaussian kernel. Prior to cortical network estimation, task structure was also regressed from the final smoothed outputs (Du et al., 2025).

### Cortical Network Estimation

Within-individual cortical networks were estimated using a Multisession Hierarchical Bayesian Model (MS-HBM) on all available task-regressed BOLD runs (Du et al., 2024; 2025; see also Kong et al., 2019). Here, we outline the key steps. For each run, correlations were computed between the timeseries of each cortical surface vertex and 1,175 uniformly distributed regions across the surface. An initial functional connectivity profile was then generated by binarizing the top 10% of these correlations. The MS-HBM was initialized with a group-level prior derived from the DU15NET-Consensus (Du et al., 2024) and used an expectation-maximization algorithm to assign all cortical surface vertices to one of 15 networks, plus a noise category, based on the functional connectivity profiles. Each participant was run through the model with a small group of previously published individuals’ data (5 participants total in the group, including *P7*-*P10* from Du et al., 2024). Finally, to ensure that network estimates were not overly influenced by model assumptions, model-free seed-region correlation checks were performed to validate all higher-order network estimates in each participant.

Two groups of association networks were examined that have previously revealed antagonistic effects: AMN (also termed the cingulo-opercular network) and SN versus DN-A and DN-B. AMN and SN respond preferentially to salient targets that are being acted upon (Seeley et al., 2007; Menon and Uddin, 2010; Seeley, 2019; Du et al., 2024; Dosenbach, Raichle, and Gordon, 2025), while DN-A and DN-B together are suppressed during such tasks (e.g., Du et al., 2024; see also Buckner and DiNicola, 2019). That is, during target detection, DN-A/DN-B both robustly oppose AMN/SN and were selected *a priori* as the focus of our analyses.

### Analysis of Task-Evoked BOLD Responses

Task-evoked BOLD responses were analyzed within the general linear model (GLM) using FSL’s first-level FEAT (v5.0.4; Jenkinson et al., 2012; Woolrich et al., 2001). Surface-projected data were high-pass filtered (0.01 Hz) and entered into a GLM with targets, lures, start/end cues, and extended fixation blocks as regressors; distractor nontargets served as the implicit baseline. The contrast of interest was targets (red Ks) versus distractor nontargets (black Ks and Os). FEAT estimated β values at each surface vertex for this contrast and converted them to z values. The resulting z-value maps were then averaged across runs using *fslmaths* to produce a single task activation map within each participant (**Fig. 1**, top three rows). Individual participant maps were also averaged to create a group-level map (**Fig. 1**, bottom row).

To examine the temporal profile of the target detection response within functional networks, trial-locked BOLD timeseries were extracted from within the borders of AMN/SN and DN-A/DN-B. For each run, the mean signal within each group of networks was epoched from 3 seconds before to 15 seconds after each target and nontarget trial onset. Each epoch was baseline-corrected by subtracting the mean of the 3-second pre-stimulus period, separately for target and nontarget trials. The difference between target and nontarget epochs was then computed at each timepoint to produce a contrast timeseries reflecting the differential BOLD response. For individual participants, the mean and standard error of the mean of this difference were computed across runs (**Fig. 2** left panel, top three rows); for the group, across participants (**Fig. 2** left panel, bottom row). Values are expressed in units of raw signal change from the pre-stimulus baseline, with positive values indicating greater activation for targets relative to nontargets.

### Intracranial Electrode Implantation and iEEG Data Acquisition

After the pre-operative fMRI session, each participant was implanted with clinical depth electrodes, in most cases the MICRODEEP Depth Electrodes (DIXI Medical, Marchaux-Chaudefontaine, France). In three participants, both MICRODEEP Depth Electrodes and Behnke-Fried Macro-Micro Depth Electrodes (Ad-Tech, Oak Creek, WI, USA) were implanted. In addition to standard macroelectrode contacts, each Behnke-Fried electrode contains a bundle of eight microwires for recording single-neuron activity, extending approximately 4 mm beyond its distal tip (Misra et al., 2014); each of these participants had three microwire bundles. Electrode trajectories were determined at a clinical case conference independent of this research protocol. Post-operatively, participants underwent a CT scan and stayed in the Epilepsy Long-Term Monitoring Unit for localization of seizure foci. Intracranial recordings were obtained using a 128-channel neural signal processor for two participants (*P1*, *P2*) and a 256-channel neural signal processor for four participants (*P3*-*P6*), sampled at 1000 Hz (Blackrock Microsystems, Salt Lake City, UT, USA). All microwire signals were sampled at 30 kHz. Electrode contacts were referenced to a contact located within the white matter (Mercier et al., 2022).

Participants were invited to complete the same target detection task as from the pre-operative fMRI session (with unique stimulus ordering). Task stimuli were presented via PsychoPy on a monitor at the bedside, and participants indicated their responses to targets by pressing the space bar on a keyboard. Prior to each run, participants were instructed to stay still and alert. No participant experienced a seizure during any data collection session. Runs were excluded if the task was interrupted by a clinical team member, if significant participant movement was observed, or if the participant was behaviorally non-compliant on the task (see Supplementary Table S2 for details).

### Localization of Electrode Contacts with Functional Networks, Thalamus and Hippocampus

To co-localize electrode contacts, surface-based network estimates from the pre-surgical fMRI session were projected from the fsaverage6 surface into each participant’s native T1w volume space via an empirical procedure detailed in Du et al. (2024). Briefly, the same spatial transformations applied to each participant’s BOLD data were used to map each surface vertex’s network label back to its corresponding native-space volumetric coordinate; the resulting sparse volume was filled via nearest-neighbor interpolation and masked with the cortical ribbon. The final volumetric estimates were projected back to the surface to confirm spatial fidelity with the original surface-based estimates. Electrode contacts were then localized within the volumetric network estimates, as described below.

First, each patient’s pre-operative T1w MRI and post-operative CT were co-registered using custom MATLAB software and SPM12 (Friston et al., 2006). Electrode contacts were then automatically detected and assigned xyz coordinates using the Localize Electrodes Graphical User Interface (LeGUI; Davis et al., 2021). These coordinates were transformed into the space of the volumetric network estimates (1 mm isotropic) using FSL *FLIRT* and *img2imgcoord* (v6.0.7.15) and assigned to the nearest voxel. Around each contact’s center voxel, three cubic neighborhoods were defined: 3×3×3 mm (±1 voxel), 5×5×5 mm (±2 voxels), and 7×7×7 mm (±3 voxels).

For functional network localization, only contacts near or within the cortical ribbon were considered. A contact was included as a candidate if at least 10% of its ±3-voxel neighborhood overlapped with the cortical ribbon and the nearest cortical ribbon voxel was less than 3 mm away. Contacts whose center voxel fell within the cortical ribbon were classified as gray matter; others were classified as white matter. Contacts localized to the noise category (60 total) were not considered candidates for analysis. A gray matter contact was assigned to AMN/SN or DN-A/DN-B if three criteria were met: (1) the center voxel overlapped with the network, (2) more than 50% of gray matter voxels in the ±1-voxel neighborhood shared the network label, and (3) more than 25% of gray matter voxels in the ±2-voxel neighborhood shared the label. A white matter contact was assigned to AMN/SN or DN-A/DN-B if three criteria were met: (1) the adjusted^1^ modal network label within the ±3-voxel neighborhood was AMN/SN or DN-A/DN-B; (2) the label agreed with the network label at the nearest gray matter voxel; and (3) the modal network comprised at least twice the proportion of the next most common network label within the neighborhood (this criterion was waived if the next most common label was “Unknown” or in the noise category).

As an additional check, seed region-based functional connectivity maps were computed for each contact. This additional validation analysis derived an estimate of the network associated with each contact without network model assumptions. For this analysis, a cubic region around each contact (±1 voxel for gray matter, ±3 voxels for white matter) was projected to the fsaverage6 cortical surface. For each run, the mean BOLD timeseries within the projected region was correlated with every other surface vertex. The resulting Fisher z-transformed correlation maps were averaged across runs and then quantified by computing the mean z(r) within the borders of each association network. Every candidate contact was visualized in axial, coronal, and sagittal views overlaid on the native T1w image and volumetric network estimates, alongside the seed-based correlation maps and all criterion values, via an automated HTML report that was carefully reviewed for each patient. Contacts with borderline criterion values, ambiguous correlation maps, or locations near network boundaries were flagged and reviewed at consensus meetings, where final network assignments were determined using all available evidence (criterion values, volumetric visualizations, and correlation maps). All assignments were finalized prior to iEEG analysis.

Thalamic and hippocampal contacts were localized in a manner similar to the cortical contacts, using the FreeSurfer anatomical segmentation (aparc+aseg) rather than network estimates. A contact was localized to the thalamus if three criteria were met: (1) the adjusted ^1^ modal anatomical label within the ±3-voxel neighborhood was “Left/Right-Thalamus”; (2) this label comprised at least 20% of the ±3-voxel neighborhood; and (3) the contact was within the five most medial contacts on the depth electrode targeting the thalamus (this criterion was waived if 40% or more of the neighborhood was “Left/Right-Thalamus”). A contact was localized to the hippocampus if two criteria were met: (1) the modal anatomical label within the ±3-voxel neighborhood was “Left/Right-Hippocampus” (if the modal label was white matter or unlabeled, the next most common label was used instead); and (2) this label comprised at least 20% of the ±3-voxel neighborhood. For contacts localized to the hippocampus, subregion labels (head or body) were assigned using the Tian Melbourne Subcortex Atlas (S4, 3T; Tian et al., 2020) in Montreal Neurological Institute (MNI) space. Electrode coordinates were transformed to MNI space (1.0 mm isotropic) and the adjusted modal Tian label within the ±3-voxel neighborhood was used for subregion assignment. Similar to cortical contacts, every candidate thalamic and hippocampal contact was visualized in an automated HTML report with axial, coronal, and sagittal views overlaid on the native T1w image, the FreeSurfer anatomical segmentation, and the Tian subcortical atlas for hippocampal contacts (Tian et al., 2020), alongside all criterion values. Each report was carefully reviewed, and contacts with borderline values or ambiguous locations were flagged and resolved at consensus meetings. All assignments were finalized prior to iEEG analysis.

Microwire bundles were putatively localized to AMN/SN or DN-A/DN-B for each participant with available single-neuron recordings by visual inspection of cortical network assignments overlaid on the pre-operative T1w MRI in LeGUI, co-registered with SPM12. Each microwire was then individually reviewed at a consensus meeting with neurosurgeons (B.S., S.R.) to determine the putative network that each microelectrode was recording from, prior to single-neuron analysis.

### iEEG Data Quality Control and Preprocessing

Raw electrophysiological recordings for every iEEG contact in every run were visually inspected for quality. Contacts that showed either flat recordings or high frequency noise throughout a task run were excluded for that run. A total of 40 contacts were excluded for this reason. Reference and ground contacts were also excluded (2 per participant, 12 total). Contacts within seizure onset zones, as determined by an attending clinical neurologist, were excluded, except those at or near the hippocampus, which were retained to preserve hippocampal sampling. All contact/run exclusions were finalized prior to additional preprocessing and analysis to avoid bias. As a further control analysis, we also repeated the analyses excluding hippocampal seizure-onset-zone contacts, which did not alter the effects observed in the hippocampus (Supplementary Fig. S4).

iEEG data were preprocessed using a custom pipeline built on MNE-Python (Gramfort et al., 2013). Line noise was removed with zero-phase Finite Impulse Response (FIR) notch filters at 60, 120, and 180 Hz (5 Hz bandwidth). Contacts were then common average re-referenced. Two copies of the preprocessed data were generated: one downsampled to 500 Hz for analysis of the high-frequency broadband (HFB) response, and one retained at 1000 Hz for time-frequency spectrograms. The first trial of each run was dropped to avoid modeling task onset transients.

### Analysis of Task-Evoked HFB Responses

We first examined HFB activity, which tracks local population firing and correlates with the BOLD signal (Logothetis et al., 2001; Mukamel et al., 2005; Nir et al., 2007; Manning et al., 2009). For each contact, HFB envelopes were computed from the preprocessed 500 Hz data as follows. The 75–175 Hz range was divided into ten, linearly spaced, 10 Hz-wide sub-bands (75–85, 85–95, …, 165–175 Hz), each isolated with a zero-phase FIR bandpass filter (firwin design). The envelope of each sub-band was extracted as the magnitude of the analytic signal obtained via the Hilbert transform, and the resulting envelopes were averaged across sub-bands to yield a single broadband envelope per contact.

HFB envelopes were lightly smoothed with a Gaussian kernel (σ ≈ 8.3 ms; 50 ms window) to reduce high-frequency noise. Solely for the purpose of visualizing HFB responses (e.g., **Figs. 2** and **3**), a wider kernel (σ ≈ 67 ms; 400 ms window) was substituted. After smoothing, envelopes were segmented into single-trial epochs spanning −1 to 2 sec relative to stimulus onset for target and nontarget trials separately, and a systematic outlier rejection procedure was applied (see *Trial-Level Outlier Rejection*). The contrast of interest was computed as the percent change of each target trial’s HFB envelope relative to the mean nontarget HFB envelope within the same run, time-resolved across the trial epoch. For each contact, contrasts were averaged across runs.

#### Trial-Level Outlier Rejection

To mitigate the effects of transient artifacts such as interictal epileptiform discharges (IEDs), systematic outlier rejection was applied across trials, after aligning data to the stimulus onsets. HFB envelopes were screened for outliers independently for each contact and run, separately for target and nontarget trials. The peak-to-peak amplitude of every trial envelope was computed, and trials exceeding 2 standard deviations from the mean were rejected. Peak-to-peak amplitude was chosen because transient high-amplitude deflections such as IEDs are better captured by dynamic range than by mean amplitude, a consideration particularly relevant for contacts within seizure onset zones. Trial rejections determined from HFB envelopes were carried forward to subsequent analyses.

#### Statistical Testing

Cluster-based permutation tests (Maris and Oostenveld, 2007) were performed to assess the statistical significance of the task contrast for each contact (**Fig. 3**). Per-run contrast timeseries were binned into non-overlapping 50 ms bins, and a one-sample t-test against zero was computed across runs at each bin. Contiguous bins exceeding an uncorrected threshold of *p* < 0.05 were grouped into temporal clusters, including single-bin clusters, with positive and negative clusters identified separately to prevent merging of activity increases and decreases. Each cluster was assigned a mass equal to the sum of absolute t-values across its bins. Analogous to the within-subjects permutation scheme in Maris and Oostenveld (2007) with runs as the unit of observation, a null distribution was generated by randomly flipping the sign of each run’s contrast timeseries (5000 permutations), recomputing binned t-statistics, and retaining the maximum cluster mass from each permutation. Family-wise error rate was controlled by comparing each observed cluster mass against this null distribution of the permutation maximum; the corrected *p*-value was the proportion of permutation maxima equaling or exceeding the observed mass. Because contacts were assigned *a priori* to specific networks and subcortical regions, family-wise correction was applied across time bins within each contact rather than across contacts. Clusters with a corrected *p* < 0.05 were considered significant. We refer to significant negative clusters (lower HFB activity for targets than nontargets) as suppression.

#### Quantitative Timing Estimates

To quantify the temporal ordering of observed HFB responses across networks and regions, we leveraged the cluster-based permutation results described above. Each network and region was assigned an expected response direction based on the observed data (e.g., **Fig. 3**): an increase for AMN/SN and thalamus, a decrease for DN-A/DN-B and anterior hippocampus (head), and a biphasic increase-then-decrease for posterior hippocampus (body). A contact was included in the primary timing analysis if its earliest significant cluster (of either sign) began between 0 and 1000 ms after stimulus onset and it had at least one significant cluster overlapping the 0 to 2000 ms window following stimulus onset whose sign matched the expected direction. For posterior hippocampal contacts, each phase (increase/rise and decrease/fall) was evaluated independently. Significant clusters opposite to the expected direction, including those of non-conforming contacts, are shown as gray markers in **Fig. 4** and did not contribute to the primary timing estimates. Two latencies were measured for each conforming contact: onset, defined as the start of the earliest significant cluster matching the expected sign, and time-to-extremum, defined as the time of the peak (for increases) or trough (for decreases) within that cluster. When a contact had multiple significant clusters of the same sign, onset was taken from the earliest cluster and the extremum was searched across the interval from the earliest cluster onset to the latest cluster offset.

For each pair of networks/regions, we cross-correlated their pooled, z-scored HFB waveforms, sliding one relative to the other across lags of ±500 ms and taking the best lag at which the correlation was strongest; a positive lag indicates that the first region leads the second. As some pairs were observed to move together (e.g., AMN/SN and thalamus) and others in opposite directions (e.g., AMN/SN and DN-A/DN-B), we examined positive correlation peaks in the former case and negative in the latter. A pair was treated as simultaneous when its best-lag value exceeded the zero-lag value by less than 0.05 (in z-scored cross-correlation units), i.e., when shifting one waveform had minimal gain over alignment at zero. Because the posterior hippocampus was biphasic, only its rising phase was cross-correlated against networks and the thalamus; for these pairs the analysis window was restricted to 0 ms through the post-peak trough of the participant’s averaged hippocampal-body waveform (Supplementary Methods). Its falling phase was omitted, and the two hippocampal regions (head and body) were not paired, as their slow, late-diverging suppression profiles are poorly suited to a ±500 ms lag analysis.

Results were pooled in two stages with equal weighting across participants. Latencies were measured per contact and summarized within each participant as the median across conforming contacts (**Fig. 4**, top three rows). Lags were computed once per participant for each region pair, from network/region-averaged waveforms. These per-participant values (median latencies and best lags) were then averaged across participants (**Fig. 4**, bottom row). Given the small sample size, timing effects are reported descriptively. Group timing values are reported as mean ± SD across participants.

### Time-Frequency Spectrograms

To characterize the full spectral profile of the target detection response, time-frequency spectrograms were computed on the preprocessed 1000 Hz data. Data were segmented into epochs spanning −1 to 3 sec relative to stimulus onset to include extra data in order to avoid wavelet edge effects, which are most pronounced at low frequencies (reaching approximately ±0.55 sec at 3 Hz). Trials flagged as outliers during HFB screening (see *Trial-Level Outlier Rejection*) were removed.

Single-trial power was estimated using Morlet wavelets at 42 logarithmically spaced frequencies from 3 to 250 Hz, with cycle counts increasing linearly from 3 to 10 across frequencies. This yielded spectral resolution (FWHM) ranging from ∼2.4 Hz at 3 Hz to ∼59 Hz at 250 Hz, and temporal resolution from ∼375 ms to ∼15 ms. Each wavelet was truncated at ±3.5 temporal standard deviations, normalized to unit energy, and convolved with each contact’s single-trial timeseries via fast Fourier transform (FFT). Instantaneous power was computed as the squared magnitude of the resulting complex coefficient and downsampled in time by a factor of 10 to 100 Hz.

For each contact and run, single-trial power was converted to amplitude and averaged within each trial type. The contrast of interest was computed as the percent change of mean target amplitude relative to mean nontarget amplitude within each run, analogous to the HFB contrast, and per-run contrasts were averaged across runs. As with the HFB analysis, all available contacts within each network (AMN/SN, DN-A/DN-B) or region (thalamus, hippocampal head, hippocampal body) were averaged within each participant (**Figs. 5** and **6**, top three rows), then across participants to yield a grand average (**Figs. 5** and **6**, bottom row).

### Analysis of Single-Neuron Firing Rates

#### Spike Sorting

Offline Sorter (Plexon, Dallas, TX) was used to isolate units within the neural data recorded from microwires. First, putative action potential waveforms were detected as any local minima exceeding a threshold of -3.5 root mean squared from each microwire. All microelectrode channels were initially analyzed using an automatic spike sorting algorithm, employing a T-Distribution Expectation Maximization sorting method. Then, the principal component space for each channel was visually inspected to remove noise detections, and detect any additional clusters not detected by the automatic sorter. Any additional clusters found through manual sorting were added to the sample.

#### Classification of Putative Cell Subtypes

To classify the units into cell subtypes, we quantified features of each action potential waveform using two metrics: FWHM and trough-to-peak duration (TTP). Two clusters were observed, representing putative inhibitory interneurons (IN) and excitatory principal cells (PC). INs were classified as units with a FWHM ≤ 0.3 ms and a TTP ≤ 0.3 ms. These cutoffs marked the clearest separation between the two clusters in this sample (**Fig. 7B**). Because no standard cutoff exists and waveform distributions differ across samples, they differ slightly from the cutoff used by Cowan et al. (2025).

#### Neuronal Firing Rate Calculations and Statistical Testing

For each isolated neuron, average firing rates were calculated from stimulus-aligned spikes in nonoverlapping 50 ms bins. For each neuron, bin counts were converted to Hz and standardized using the mean and standard deviation of firing across all target, nontarget, and lure baseline bins from −0.5 to −0.1 sec: zFR = (bin firing rate − baseline mean) / baseline SD. The population permutation test calculated each neuron’s FR difference between target and nontarget trials at each time bin between 0 to 2 sec, randomly reversed the sign of those paired differences across neurons 1000 times, and compared the observed mean difference with the two-sided permutation distribution at α = 0.05. The displayed condition means and SEM were Gaussian-smoothed over 100 ms (**Fig. 7E**, **7F**).

## Acknowledgements

We thank the Harvard FAS Division of Research Computing. We thank Tim O’Keefe and Daniel Asay for assisting in optimization of data processing, Ross Mair for MRI physics support, and Anne Billot, Xiangyu Wei, Mark Eldaief, and Vaibhav Tripathi for helpful feedback and discussion.

## Footnotes

1 For candidate cortical contacts, if the modal label was “Unknown” i.e., unassigned, the next most common label was used instead. For candidate thalamic or hippocampal contacts, if the modal label was “Unknown” i.e., unassigned, or “Left/Right-Cerebral-White-Matter”, the next most common label was used instead.

